# Reinforcing Nicotine Doses Activate Supramammillary VGluT2 Neurons in Mice

**DOI:** 10.64898/2026.08.13.744511

**Authors:** Yosuke Arima, Xia Min, Beyonce Getachew, Laura D. Nicolas, Alex Gillespie, Ana Armenta Vega, Sarah T. Johnson, Guohua Bi, Zengyou Ye, Satoshi Ikemoto

## Abstract

**Background:** Although nicotine reinforcement is often attributed to mesolimbic dopamine neurons in the ventral tegmental area, accumulating evidence indicates that additional brain circuits contribute to its reinforcing effects.

**Aims:** The hypothalamic supramammillary region (SuM) has been implicated as one such substrate, yet the cellular targets and circuit mechanisms through which nicotine engages this region remain poorly understood.

**Methods:** We combined RNAscope in situ hybridization to identify nicotinic acetylcholine receptor (nAChR) subunits, intravenous nicotine self-administration in mice to determine doses that reliably support reinforcement, and fiber photometry to monitor calcium activity in SuM VGluT2 neurons in vivo.

**Results:** Mice exhibited reliable nicotine self-administration across a range of doses under fixed-ratio and progressive-ratio schedules. RNAscope analysis revealed prominent expression of the β2 nAChR subunit in VGluT2-expressing neurons projecting from the SuM to the medial septum. Fiber photometry recordings showed that reinforcing doses of nicotine produced rapid, infusion-locked increases in GCaMP signals in SuM VGluT2 neurons.

**Conclusions:** These findings identify nAChR-expressing SuM neurons as a candidate circuit substrate engaged by reinforcing doses of nicotine and extend current models of nicotine reinforcement beyond canonical mesolimbic dopamine pathways.

## INTRODUCTION

The mesolimbic dopamine system has long been implicated in the reinforcing effects of nicotine (Corrigall et al., 1992), yet accumulating evidence indicates that additional neural circuits contribute to nicotine reinforcement (Wills et al., 2022). Notably, rats will self-administer nicotine directly into the supramammillary region (SuM) (Ikemoto et al., 2006), indicating that nicotine can act locally within this hypothalamic structure to support reinforcement. Despite this observation, the cellular targets and circuit mechanisms through which nicotine engages SuM neurons remain poorly understood.

Recent work has further implicated the SuM in reinforcement-related processes. Optogenetic activation of SuM vesicular glutamate transporter 2 (VGluT2)-expressing neurons (SuM^VGluT2), including those projecting to the medial septum (SuM^VGluT2→MS), supports reinforcement and elicits dopamine release in the nucleus accumbens (Kesner et al., 2021). These findings raise the possibility that nicotine directly engages SuM circuits via nicotinic acetylcholine receptors (nAChRs) expressed by SuM neurons. However, the expression of nAChRs subunits within defined SuM neuronal populations has not been systematically examined.

Interpretation of nicotine reinforcement mechanisms also depends on behavioral paradigms that reliably index reinforcing efficacy (Thomsen et al., 2009; Fowler and Kenny, 2011). Intravenous self-administration (IVSA) remains the gold standard for assessing drug reinforcement, yet validated nicotine IVSA procedures in mice are limited. Establishing dose ranges that consistently support nicotine self-administration in mice is therefore essential for circuit-level analyses.

Here, we addressed these gaps by (1) characterizing nicotinic acetylcholine receptor subunit expression in SuM neurons, with particular emphasis on SuM^VGluT2→MS projection neurons; (2) evaluating intravenous nicotine self-administration under fixed-ratio and progressive-ratio schedules to define reinforcing dose ranges in mice; and (3) determining whether reinforcing doses of nicotine activate SuM VGluT2 neurons in vivo using fiber photometry. Together, these experiments integrate molecular, behavioral, and physiological approaches to identify SuM VGluT2 neurons as a candidate substrate for nicotine reinforcement.

## MATERIALS AND METHODS

### Animals

C57BL/6J mice were obtained from The Jackson Laboratory. VGluT2-Cre mice (Slc17a6^tm2(cre)Lowl/J^) on a C57BL/6J background were bred at the National Institute on Drug Abuse (NIDA). Mice were housed under a reversed light–dark cycle with ad libitum access to food and water, except during IVSA experiments, in which body weights were maintained at 85–90% of baseline. Food restriction is recognized for amplifying the reinforcing effects of substances (Cabeza de Vaca and Carr, 1998) and is commonly used in nicotine IVSA procedures with rodents (Contet et al., 2010; Fowler and Kenny, 2011). This degree of food restriction in rodents has been shown to improve their health, rather than cause harm (Speakman and Mitchell, 2011). All procedures were approved by the NIDA Animal Care and User Committee and conducted in accordance with institutional guidelines.

### Drugs

Nicotine ditartrate dihydrate (Sigma-Aldrich) and nicotine pyrrolidine methiodide (NPM) were prepared based on free-base concentrations and dissolved in sterile saline. The pH of the solution was adjusted to ∼7.0 using NaOH. NPM, cocaine hydrochloride, and fentanyl citrate were obtained from the NIDA Drug Supply Program. Either methohexital (Brevital; 1%, 0.02–0.03 mL) or propofol (20 mg/kg) were used to verify catheter patency when catheter failure was suspected.

### Surgical Procedures

Intra-medial septum (MS) injections: For RNAscope experiments, mice were anesthetized with isoflurane (1–2%) and placed in a stereotaxic frame. Cholera toxin subunit B (CTb; 0.5%, 100 nL) was injected into the medial septum (MS).

Jugular vein catheter implantation: Mice were anesthetized with ketamine/xylazine and implanted with a chronic indwelling jugular vein catheter. Catheters were flushed daily with 0.1 mL of heparin/gentamicin solution to maintain patency. After 3 days of recovery, food restriction was resumed.

Intra-supramammillary region (SuM) surgery: For fiber-photometry experiments, mice first received jugular catheter implantation and subsequently underwent stereotaxic surgery for viral delivery. AAV9-pGP-AAV-syn-FLEX-jGCaMP7s-WPRE (Addgene; titer 1.0 × 10^13^ gc/mL) was injected into the SuM. An optic fiber was implanted with the fiber tip positioned at the SuM and secured to the skull with dental cement.

### RNAscope In Situ Hybridization

Tissue preparation: Two-month-old male and female C57BL/6J mice were deeply anesthetized with isoflurane and transcardially perfused with 10% formaldehyde in 0.1 M phosphate buffer (pH 7.3). Brains were post-fixed for 2 h, cryoprotected sequentially in 20% and 30% sucrose solutions at 4°C, and coronally sectioned at 14 μm.

RNAscope procedure: RNAscope assays were performed according to the manufacturer’s instructions using probe sets targeting nicotinic acetylcholine receptor subunits α3, α4, α7, β2, and β4, as well as VGluT2, VGAT, GAD65, and GAD67. Sections hybridized with the bacterial DapB probe served as negative controls and showed no detectable signal.

Imaging and analysis: Sections were imaged using an Olympus FV1000 confocal microscope. RNAscope images were analyzed using FIJI and QuPath. Cell detection was performed based on nuclear staining with DAPI or CTb, followed by subcellular spot detection on the RNA channel. Detection parameters were adjusted conservatively for each image to minimize false-positive detection of RNA puncta, while downstream classification was performed using fixed, objective criteria. For each detected cell, the number of RNA puncta (spot count) and the total area of RNA clusters were extracted from QuPath measurement tables. RNA-positive cells were classified using a combined criterion based on puncta count or total cluster area, independent of fluorescence intensity. Cells were assigned to hierarchical positivity levels according to the following thresholds: ≥2 puncta or ≥0.4 µm².

Regions of interest (ROIs) corresponding to the SuM were manually delineated based on anatomical landmarks. For each brain, SuM ROIs were defined at anterior, middle, and posterior levels, and cells within each ROI were quantified separately. Analyses were performed using samples obtained from four male and four female mice.

### Operant Conditioning Chambers

Experiments were conducted in standard operant conditioning chambers (Med Associates) equipped with two levers, cue lights, and a house light. One lever was designated active and the other inactive. Responses on the active lever resulted in programmed reinforcement; inactive lever presses had no consequence. Cue lights above the active lever were illuminated during nicotine infusions but not during food self-administration. Food magazines were present only during food-training sessions.

### Intravenous Self-Administration (IVSA) Schedules

Two reinforcement schedules were used: a fixed-ratio 5 schedule with a 20-s timeout (FR5) and a modified progressive-ratio 3 schedule (PR3), in which the response requirement increased by three responses after each reinforcement. Prior to nicotine testing, mice were trained to respond for food pellets (20 mg; Bio-Serv).

#### FR5 Nicotine Self-Administration

Mice were trained to self-administer food pellets for 7 consecutive days prior to catheter implantation. Training began on an FR1 schedule and progressed to FR5 during 60-min sessions. After training, mice were returned to ad libitum feeding for 24 h before surgery.

Following catheter implantation and recovery, VGluT2-Cre mice were retrained for 8 – 10 days under the FR5 schedule with food reinforcement. Nicotine self-administration procedures were adapted from Fowler and Kenny (2011). Mice then self-administered nicotine (30 µg/kg/infusion; 35 µL delivered over 3 s) paired with cue-light illumination during 60-min sessions for 15 days.

Dose–response testing was subsequently conducted using nicotine doses of 30, 100, 0, and 10 µg/kg/infusion presented sequentially. Mice were required to demonstrate stable intake (<20% variation across three sessions) before advancing to the next dose. Although intravenous surgery was conducted on 12 mice (6 males, 6 females), catheter failure resulted in only 6 mice (3 males, 3 females) completed the protocol and were included in analyses.

#### Modified PR3 Nicotine Self-Administration

To ensure that VGluT2-Cre and C57BL/6J mice respond to nicotine in a comparable manner, both strains were used. This is crucial for the fiber-photometry experiment outlined below. Mice were first trained to respond for food pellets under an FR1 schedule prior to catheter implantation. Sessions lasted 60 min. After surgery and recovery, mice were randomly assigned to nicotine or saline groups.

Self-administration was conducted under a modified progressive-ratio schedule in which the response requirement increased by three responses after each infusion (PR3). Unlike conventional PR schedules, no breakpoint was imposed; instead, sessions ended after 90 min regardless of response output. Because the modified PR3 schedule did not include a breakpoint criterion, session duration rather than breakpoint was used as the limiting factor for responding. This design was adopted to allow comparison of response output across nicotine doses while minimizing session termination due to early breakpoint attainment.

Mice were initially trained under a PR1 schedule with either 25 µg/kg nicotine or saline (35 µL delivered over 3 s) paired with cue-light illumination (3 s) for 7 sessions. The nicotine group were then exposed to 50 and 25 µg/kg nicotine under the PR1 and PR3 schedules. Dose– response testing was conducted in descending order (100, 50, 25, and 0 µg/kg/infusion), with each dose tested for three sessions. Saline controls received saline infusions throughout the experiment.

To reduce losses from catheter failure, each dose was tested over three sessions. This improved retention compared to the FR5 experiment: Of 33 mice that underwent surgery, 24 mice (14 nicotine, 10 saline) completed the experiment, and their data were analyzed.

#### Data Analysis

Response levels varied across sessions, particularly under the PR3 schedule. Therefore, median infusion numbers and lever responses across the final three sessions of each dose were used for statistical analyses. Because the modified PR3 schedule did not include a breakpoint criterion, reinforcement strength was assessed using the number of infusions earned and total active lever responses during the session. These measures provide a continuous index of motivation under time-limited PR procedures.

### Fiber-Photometry Recording

Recordings were performed 3–5 weeks post-surgery using a Doric fiber-photometry system. Excitation wavelengths were 465 nm (GCaMP) and 405 nm (isosbestic control). Fluorescence signals were acquired at 1,200 Hz and synchronized with infusion events.

#### Passive drug administration

Mice were habituated to the test chambers for 3 days prior to recording. Six infusions (35 µL/infusion over 3 s) of a single dose were delivered at 5-min intervals for each session. Sessions included saline, nicotine (10, 30, 100 µg/kg/infusion), cocaine (300 µg/kg), or fentanyl (6 µg/kg), and these nicotine doses and the other drugs were tested in this order with at least 24 h between sessions. Effects of nicotine pyrrolidine methiodide (NPM; 30 and 56 µg/kg/infusion) were evaluated in a separate group of mice, tested for NPM, nicotine, and saline, delivered at 15-min intervals.

#### Histology

After experiments, mice that underwent intracranial surgery were perfused with PBS followed by 10% formalin. Brains were cryoprotected, sectioned at 40 µm, and mounted with ProLong Diamond. AAV expression and fiber placement were verified using a Keyence BZ-X710 fluorescence microscope.

#### Fiber-Photometry Data Analysis

Fiber-photometry data were analyzed using custom Python scripts. Fluorescence signals from the GCaMP (465 nm) and isosbestic reference (405 nm) channels were smoothed using a zero-phase moving average low-pass filter. To correct motion artifacts and photobleaching, the isosbestic signal was linearly fitted to the GCaMP signal to generate a fitted control, and the normalized fluorescence (ΔF/F) was calculated as:

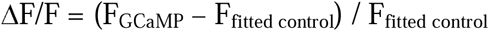

For each infusion trial, ΔF/F traces from 30 s before to 90 s after infusion onset were extracted and normalized by z-scoring using the pre-infusion period as the baseline. The area under the curve (AUC) was calculated from the mean z-score across the whole session, using a 30 s window right before and after the infusion onset. To compare effects of different IV infusion treatments on GCaMP signals, response magnitude was derived by subtracting the 30-s AUC prior to the infusion onset from the 30-s AUC post infusion onset.

## RESULTS

### Nicotinic acetylcholine receptor subunit expression in SuM**→**MS projection neurons

Nicotine delivered directly into the SuM is reinforcing (Ikemoto et al., 2006), suggesting that SuM neurons may express nAChRs. To identify potential cellular substrates, we examined the expression of nAChR subunits in SuM neurons, with a focus on SuM neurons projecting to the MS.

SuM^VGluT2→MS neurons were retrogradely labeled by CTb injection into the MS (Fig. 1A). CTb labeling was largely restricted to the SuM, with minimal labeling in adjacent regions (Fig. 1B). RNAscope in situ hybridization was used to detect mRNAs encoding nAChR subunits and neurotransmitter markers.

**Figure 1.**
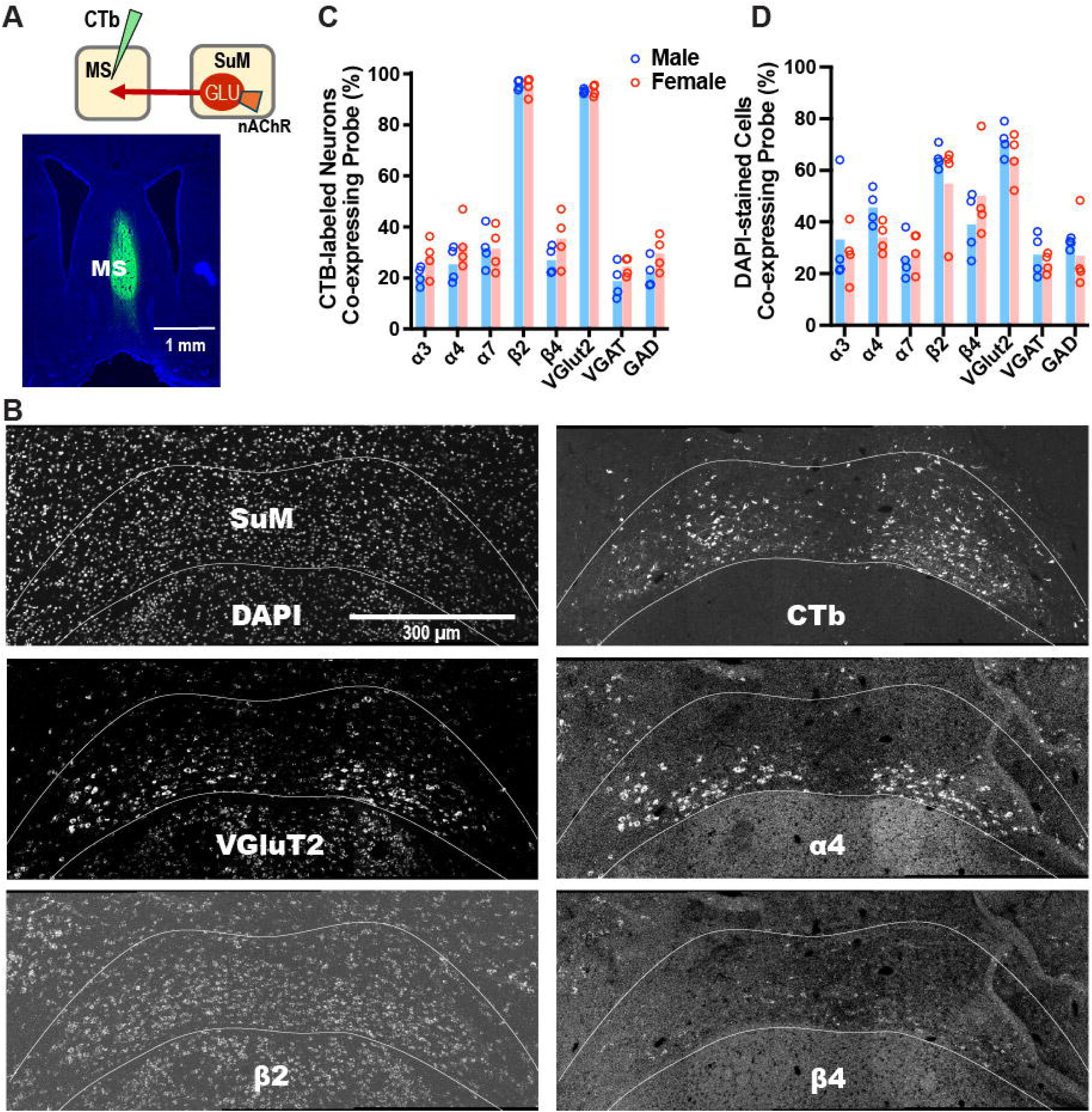
Nicotinic acetylcholine receptor subunit expression in supramammillary (SuM) neurons projecting to the medial septum (MS). (A) SuM→MS neurons were retrogradely labeled by cholera toxin B (CTb) injection into the MS, and RNAscope *in situ* hybridization was used to detect mRNAs encoding nicotinic acetylcholine receptor (nAChR) subunits and neurotransmitter markers in the SuM. (B) Representative photomicrograms illustrating mRNA probe expressions in the SuM. The images were adjusted to enhance the visibility of probe expressions at this magnification. (C) CTb-labeled neurons co-expressing nAChR subunits and neurotransmitter markers. (D) DAPI-labeled neurons co-expressing nAChR subunits and neurotransmitter markers.

Nearly all CTb-labeled neurons (97%) expressed the β2 nicotinic acetylcholine receptor (nAChR) subunit, and β2 expression was broadly distributed across the SuM, without obvious confinement to either the medial or lateral regions. Because almost all CTb-labeled neurons (93%) expressed vesicular glutamate transporter 2 (VGluT2), the majority of SuM→MS projection neurons are glutamatergic and express β2-containing nAChRs (Fig. 1C). Notably, 22-26% of CTb-labeled neurons expressed VGAT and GAD, suggesting that some of SuM→MS projection neurons use both GABA and glutamate for neurotransmission. In contrast, smaller proportions of CTb-labeled neurons (22-32%) expressed other nAChR subunits, including α3, α4, α7, and β4.

In the overall SuM neuronal population identified by DAPI staining, VGluT2 and β2 expression levels were lower (approximately 70% and 60%, respectively), whereas α4 and β4 expression were slightly higher (approximately 40% and 50%, respectively) (Fig. 1D). Expression of the remaining nAChR subunits was comparable between projection-defined and total SuM populations. No sex differences were observed for any nAChR subunit.

### Fixed-ratio responding reveals dose-dependent regulation of nicotine intake

To characterize intravenous nicotine self-administration (IVSA) in mice, we first evaluated operant responding under a fixed-ratio 5 (FR5) schedule (Fig. 2A). Mice self-administered nicotine at doses of 10, 30, or 100 µg/kg/infusion, or saline vehicle.

**Figure 2.**
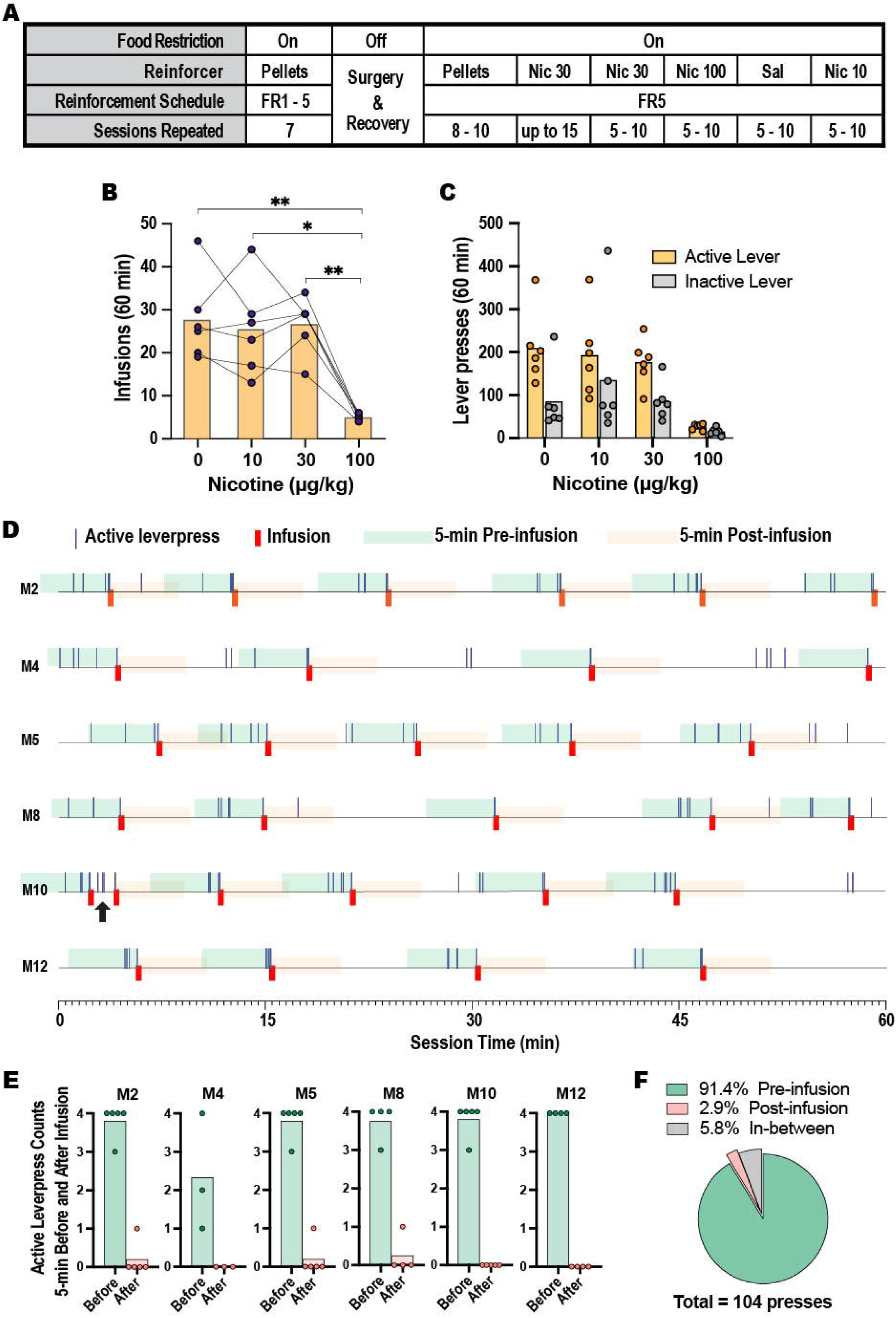
Nicotine self-administration under an FR5 schedule. (A) Timeline describing experimental events occurred (from left to right). Abbreviations: Nic#, nicotine and its dose; Sal, saline. (B,C) Number of infusions (B) and active and inactive lever presses (C) during self-administration of nicotine (0, 10, 30, and 100 µg/kg/infusion). Dots indicate individual mouse responses; bars indicate means (*n* = 6). (D) Event records showing active lever presses and infusion timing during 60-min sessions. Colored segments denote 5-min periods before and after each infusion; overlapping periods (shown by the arrow for M10) were excluded. Final infusion periods with incomplete post-infusion intervals (M2, M4, and M8) were excluded. (E) Active lever presses during 5-min pre– and post-infusion periods. The infusion-triggering response was excluded, yielding a maximum of four responses per period. (F) Probability of active lever pressing during pre-infusion, post-infusion, and inter-infusion periods, calculated from pooled data across mice.

Under the FR5 schedule, mice responded on the levers and obtained infusions across conditions (Fig. 2B,C). The highest dose tested (100 µg/kg/infusion) produced significantly fewer infusions and lever presses than the other doses (infusions: F(3,15) = 17.05, P < 0.001; lever presses: F(3,15) = 12.77, P < 0.001; Supplementary Table 1). No significant lever-by-dose interaction was detected (F(1.0,5.3) = 1.94, P = 0.221). Across all doses, mice responded more on the active lever than on the inactive lever (F(1,5) = 12.71, P = 0.016).

Event records revealed that responding at the 100-µg/kg dose exhibited a distinct temporal pattern characterized by post-infusion pauses (Fig. 2D). The mice responded primarily during the 5-min period preceding each infusion, with little responding during the 5-min period following the infusion. To quantify this pattern across all six mice, active lever responses were compared during 5-min intervals before and after each infusion (Fig. 2E). Across animals, the majority of responses occurred during the 5-min interval preceding infusions (91% of all lever presses), whereas few responses occurred during the 5-min interval immediately following infusions (3% of all lever presses) (Fig. 2F). Thus, under the FR5 schedule, the 100-µg/kg dose produced reduced response rates accompanied by pronounced post-infusion pauses.

In contrast, infusion numbers and lever responses at the 10 and 30 µg/kg doses did not exceed those observed with saline. Similar findings have been reported previously in mice responding under FR schedules (Contet et al., 2010). Studies in rats suggest that progressive-ratio schedules are more sensitive than fixed-ratio schedules for detecting reinforcing effects of moderate nicotine doses (Donny et al., 1999; Sorge and Clarke, 2011). Therefore, we next evaluated nicotine self-administration in mice using a progressive-ratio schedule.

### Progressive-ratio responding reveals an inverted U-shaped nicotine dose–response function

Nicotine IVSA was subsequently examined using a modified progressive-ratio (PR3) schedule (Fig. 3A). Because relatively high levels of responding for saline were observed under the FR5 schedule, a saline control group was included that received saline infusions throughout the experiment. Both wild-type C57BL/6 mice (n = 10) and VGluT2-Cre mice (n = 14) were tested.

**Figure 3.**
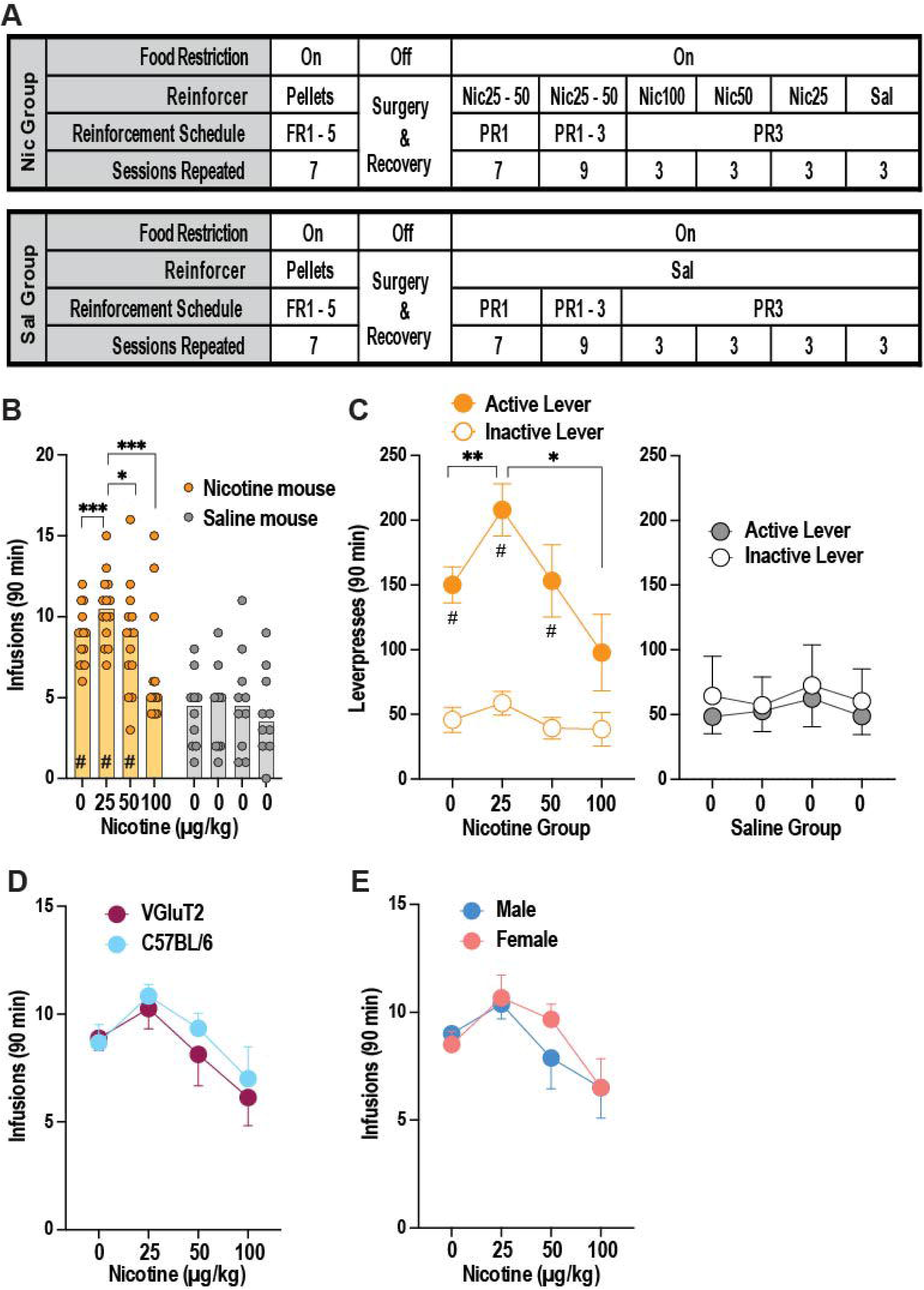
Nicotine self-administration under a PR3 schedule. (A) Timeline describing experimental events occurred (from left to right) in nicotine (Nic) and saline (Sal) groups. (B,C) Infusions (B) and lever presses (C) of 0, 25, 50, and 100 µg/kg in the nicotine group (*n* = 14) and during corresponding saline infusions of the control group (*n* = 10). Dots indicate individual mouse responses; bars indicate means. \**P* < 0.05, \*\*\**P* < 0.001; #*P* < 0.01 vs. corresponding values of the saline group (Bonferroni-corrected P values). (D) Strain effects on infusions across doses in VGluT2-Cre (*n* = 8) and C57BL/6 mice (*n* = 6). (E) Sex effects on infusions across doses in male (*n* = 8) and female (*n* = 6) mice.

Mice were assigned to either a nicotine group (n = 14) or a saline control group (n = 10). Under the PR3 schedule, nicotine self-administration exhibited an inverted U-shaped dose–response function (Fig. 3B). Infusion numbers in the nicotine group varied as a function of dose, whereas responding in the saline group remained relatively stable across sessions. Mixed ANOVA revealed a significant group-by-dose interaction (F(1.7,36.7) = 4.70, P = 0.020). Infusions obtained at the 0, 25, and 50 µg/kg doses were significantly greater than those obtained by the saline group during the corresponding sessions (Supplementary Table 1), whereas infusions at the 100 µg/kg dose did not differ from saline. Within the nicotine group, multiple comparisons showed that the 25 µg/kg dose produced significantly more infusions than the 0, 50, or 100 µg/kg doses, and the 100 µg/kg dose produced fewer infusions than the 25 and 50 µg/kg doses. Moreover, a within-subjects contract for group x dose revealed a significant quadratic trend for group x dose, F(1,22) = 9.57, P = 0.005, confirming an inverted U-shaped effect of dose in intake for the nicotine group (Supplementary Table 1).

Active and inactive lever responses were analyzed separately for the nicotine and saline groups using lever-by-dose within-subjects ANOVAs. In the nicotine group, a significant lever-by-dose interaction was observed (F(1.5,19.6) = 4.15, P < 0.05), reflecting dose-dependent changes in responding on the active lever, whereas inactive lever responding remained relatively constant across doses (Fig. 3C). In contrast, the saline control group showed no dose-related changes in responding on either lever during the corresponding sessions.

Potential effects of strain and sex on nicotine IVSA were also examined (Supplementary Table 2). Strain-by-dose and sex-by-dose mixed ANOVAs conducted on infusion numbers revealed no significant main effects or interactions for strain (main effect: F(1,12) = 0.28, P = 0.61; strain × dose: F(1.5,18.2) = 0.27, P = 0.708; Fig. 3D) or sex (main effect: F(1,12) = 0.11, P = 0.74; sex × dose: F(1.4,17.0) = 0.74, P = 0.449; Fig. 3E).

### IV administration of nicotine elicits rapid, infusion-locked activation of SuM VgluT2 neurons

IVSA experiments indicated that nicotine doses between 25 and 100 µg/kg reliably supported reinforcement in mice. We therefore tested whether nicotine within this dose range activates SuM VGluT2 neurons using fiber photometry with GCaMP (Fig. 4A). We also examined the effects of cocaine and fentanyl on SuM GCaMP signals to assess the selective response of SuM VGluT2 neurons to nicotine. When intravenous infusions are carried out by the experimenter rather than the subject, it remains uncertain whether the infused nicotine produces the same positive reinforcing effect. Nonetheless, this method is regarded as useful for gaining insight into the neural correlates activated by addictive substances, especially as it has been extensively used in research related to the dopamine system (Imperato et al., 1986; Di Chiara and Imperato, 1988; Pontieri et al., 1996).

**Figure 4.**
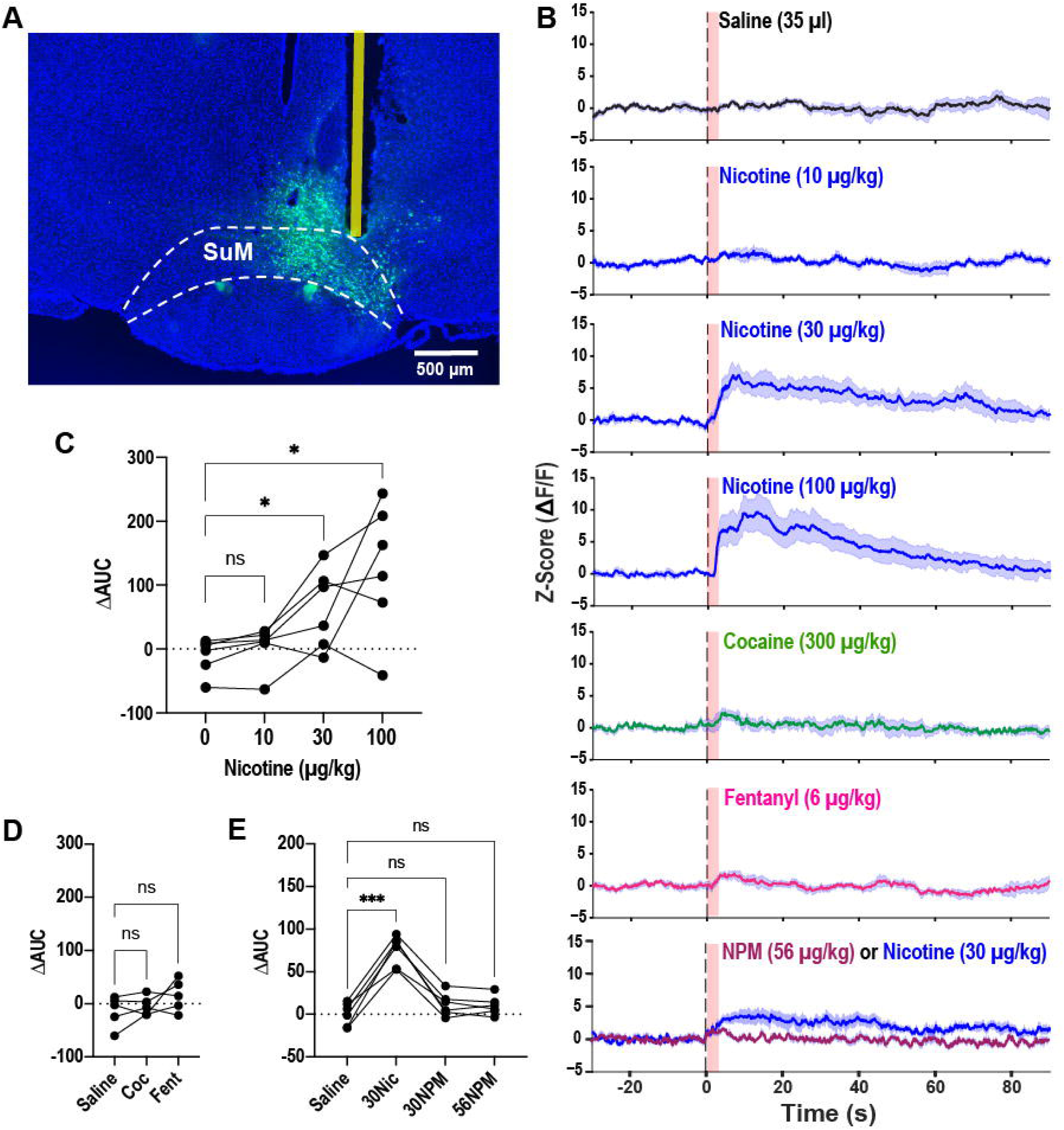
SuM GCaMP responses to intravenous drug infusions. (A) Representative GCaMP expression and fiber placement in the SuM. The photomicrogram shows fiber placement in relation to the SuM. Note that weak GFP expression just below the fiber-track tip, likely due to photo-bleaching. (B) Z-scored GCaMP signals (ΔF/F; mean ± SEM) during saline, nicotine, cocaine, fentanyl, and NPM infusions. For each drug, infusions were repeated 6 times, and each infusion (35 µl) was delivered over 3 s. The pink overlays starting at 0 on the x-axis indicate the 3-s duration of IV infusion. (C,D,E) Differences in area under the curve (AUC) for 30-s periods before vs. after infusion onset. Dots indicate individual mice. Nicotine: *n* = 6, \**P* < 0.05 (C); cocaine (COC) and fentanyl (Fent): *n* = 5 (D); nicotine (30 µg/kg) and NPM (30 or 56 µg/kg): *n* = 6, \*\*\**P* < 0.001 (E).

Intravenous nicotine produced rapid, dose-dependent increases in SuM GCaMP signals that were tightly time-locked to infusion (Fig. 4B). A one-way within-subjects ANOVA revealed a significant main effect of dose (F(3,15) = 9.34, P < 0.001; Fig. 4C). Post hoc Dunnett’s tests showed that 30 and 100 µg/kg nicotine significantly increased the GCaMP signal area under the curve (AUC) relative to vehicle. In contrast, cocaine (300 µg/kg) and fentanyl (6 µg/kg), reinforcing doses established in prior IVSA studies (Thomsen and Caine, 2006; Leonardo et al., 2023), did not reliably alter SuM GCaMP signals (F(2,8) = 2.78, P = 0.121; Fig. 4D).

Previous studies have reported that NPM, a peripherally acting nicotinic receptor agonist, alters cortical and ventral tegmental area local field potentials (Lenoir and Kiyatkin, 2011) and induces widespread c-Fos expression, including in the SuM (Dehkordi et al., 2015). To determine whether peripheral nicotinic receptor activation contributes to the SuM GCaMP response, we intravenously infused NPM and recorded SuM activity. Again, infusions of the 30µg/kg dose of nicotine increased GCaMP signals in the SuM, replicating the above experiment, whereas NPM did not significantly increase SuM GCaMP signals (Fig. 4E).

## DISCUSSION

### Nicotinic receptor expression establishes the cellular basis for nicotine activity within the SuM

This study demonstrates that the supramammillary region (SuM) exhibits nicotinic receptor expression, particularly β2-containing subunits, with notable presence in VGluT2 neurons projecting to the medial septum. The significant expression of the β2 subunit is important, given its involvement in nicotine reinforcement and self-administration across various species (Picciotto et al., 1998; Walters et al., 2006; Picciotto and Kenny, 2021). The enrichment of β2 within projection-specific SuM^VGluT2→MS neurons indicates that these cells may respond directly to nicotine and transmit nicotine-associated signals to downstream septal networks, as evidenced by behavioral and physiological analyses. Although research on nicotine reinforcement has primarily focused on the VTA and NAc, these findings underscore the importance of further investigating the contribution of the SuM to nicotine reinforcement mechanisms.

### Behavioral validation of nicotine reinforcement in mice

Nicotine intravenous self-administration in mice presents significant methodological challenges. Achieving the classic inverted U-shaped dose–response curve for nicotine is much harder than for drugs like cocaine. Even rats, commonly used for drug self-administration studies, often fail to show clear dose-dependent responses to nicotine compared to psychostimulants (Rose and Corrigall, 1997; Sorge and Clarke, 2011). These challenges are even greater in mice. For instance, Contet et al. (2010) found that mice on an FR3 schedule self-administered nicotine doses of 10–60 µg/kg/infusion at rates similar to saline and that doses of 1 and 30 µg/kg under an FR1 schedule produced comparable self-administration levels to saline, with significant reductions at 100 µg/kg. Our FR5 experiment, using 10, 30, and 100 µg/kg doses, mirrored the pattern seen by Contet et al. and similarly did not reveal an inverted U-shaped dose–response curve under fixed-ratio conditions. Conversely, Fowler and Kenny (2011), administering 30–400 µg/kg/infusion under an FR5 schedule, observed that doses between 30 and 250 µg/kg resulted in more infusions and lever presses than saline, peaking at 100 µg/kg. Although the reasons for these inconsistencies remain unresolved, their study advocated for testing higher, not lower, doses.

The absence of robust dose-response effects for nicotine must be considered alongside several important factors, such as previous self-administration exposure, the inherent reinforcing properties of cue lights, and nicotine’s dependence on contingent cues. Mice previously exposed to nicotine reinforcement displayed increased responses for saline infusions compared to controls, indicating nicotine-associated cues acquired conditioned reinforcing properties that could sustain operant behavior in its absence. Cue lights themselves can reinforce responding; both rats and mice will respond to light cues even though they have not been paired with primary rewards (Keller et al., 2014; Kish, 1955; Stewart & Hurwitz, 1958). Thus, pairing saline with a cue light can lead to substantial lever pressing. Finally, nicotine’s reinforcing effects depend heavily on contingent cue lights. Research shows intravenous nicotine self-administration is difficult to establish in rats without such cues (Caggiula et al., 2001; Donny et al., 2003), suggesting nicotine acts more as a reinforcement enhancer rather than a primary reinforcer. Altogether, these factors complicate the detection of dose-dependent responses to nicotine.

Despite this limitation, the FR5 experiment provides clear evidence that mice regulate nicotine intake. The highest nicotine dose tested reduced both infusion numbers and lever presses; however, this reduction did not reflect behavioral suppression or avoidance of nicotine. Instead, mice continued to self-administer the drug while exhibiting pronounced post-infusion pauses.

Such pauses are widely interpreted as a hallmark of regulated drug intake, reflecting titration of brain nicotine levels rather than loss of reinforcing efficacy. These findings therefore indicate that the nicotine doses used in the present study were behaviorally active and capable of supporting reinforcement in mice.

Progressive-ratio (PR) schedules are typically more effective than fixed-ratio schedules at revealing dose-dependent reinforcing effects of nicotine (Donny et al., 1999; Sorge and Clarke, 2011). Supporting this perspective, our PR testing demonstrated a distinct inverted U-shaped dose–response curve, with peak responses occurring at 25 µg/kg nicotine dose. Notably, saline control animals maintained stable behavior across sessions, suggesting that the observed changes in the nicotine group were attributable specifically to nicotine reinforcement rather than general operant activity. It is important to mention that PR schedules may be more adept at measuring the motivational aspects of drugs compared to their positive affective properties when contrasted with FR schedules (McGregor and Roberts, 1993; Donny et al., 1999). If so, nicotine may possess relatively greater motivational effects than positive affective effects relative to other addictive substances. Altogether, these behavioral results suggest that the nicotine doses used in these experiments activate reinforcement mechanisms in mice and provide essential behavioral context for interpreting SuM^VGluT2 neuronal responses to nicotine.

The mice used for intravenous self-administration were subject to food restriction. It is well established that limiting food intake affects numerous behaviors, such as substance self-administration (Carroll et al., 1979; Carr, 2007), as well as brain activity—including the dopamine system. One possible explanation for enhanced self-administration is that food restriction sensitizes the dopamine system (Carr et al., 2003; Abizaid et al., 2006; Lindblom et al., 2006; Carr, 2007; Skibicka et al., 2011) and also SuM neurons (Le May et al., 2019; Plaisier et al., 2020). Consequently, it is plausible that nicotine self-administration was partly augmented by increased SuM activity resulting from food restriction.

While our experiments did not reveal any sex differences, they were not specifically designed with sufficient statistical power to detect such differences. This limitation falls outside the scope of our study and should be considered in future research. Many rat studies have reported sex differences (e.g., (Donny et al., 2000)), though not consistently. In contrast, information about sex differences in mouse IV self-administration is far less available, and these differences may vary depending on multiple factors, such as reinforcement schedule, associated cues, strain, and administration route (Perkins et al., 1999; Pogun et al., 2017).

### Nicotine triggers rapid, infusion-locked activation of SuM VGluT2 neurons in vivo

Fiber photometry recordings revealed that intravenous nicotine induces rapid, infusion-locked increases in calcium activity among SuM VGluT2 neurons. While these experiments do not confirm whether nicotine acts directly on nicotinic receptors expressed by SuM neurons or indirectly via upstream inputs, the temporal precision of responses strongly suggests recruitment of SuM neurons following nicotine infusion. Notably, the doses eliciting robust neural responses closely correspond to those supporting reinforcement in behavioral tests.

Additionally, the absence of similar responses following cocaine, fentanyl, or a peripherally restricted nicotinic agonist indicates that SuM activation is not a general consequence of drug infusion or peripheral actions of nicotine. However, since these tests were not designed to fully evaluate these substances, comprehensive dose ranges should be explored in future research.

It is important to acknowledge that the SuM’s proximity to the VTA presents potential concerns regarding the specificity of our nicotine-induced GCaMP signal recordings within the SuM. However, previous research (Ikemoto et al., 2006) demonstrated that nicotine self-administration was associated with the SuM and posterior VTA, but not the anterior VTA, thereby mitigating these concerns. Histological analysis further confirmed that all recording sites were confined exclusively to the SuM.

Research shows that SuM neurons have nicotinic receptors and that nicotine doses, which support self-administration, activate these neurons in an infusion-specific way. This suggests nicotine triggers SuM circuits within the networks responsible for reinforcement. Previous studies found that stimulating SuM VGluT2 neurons and their pathways to the medial septum (MS) can activate MS VGluT2 neurons, which connect to the VTA and then stimulate dopamine neurons projecting to the nucleus accumbens (Kesner et al., 2021). Our current results introduce a model where nicotine activates nicotinic receptors in the SuM, stimulates SuM VGluT2 neurons, and influences mesolimbic dopamine activity through hypothalamic–septal pathways. Therefore, nicotine’s effect in the SuM may further enhance its action in the VTA by promoting dopamine neurons projecting to the NAc for nicotine’s reinforcing effects.

### Limitations and future directions

Several limitations warrant consideration. The present fiber-photometry experiments provide region-wide activity measurements and cannot determine whether nicotine preferentially activates specific SuM VGluT2 projection-defined ensembles; future studies employing projection-specific calcium imaging are needed. Additionally, selective inhibition of specific projections during nicotine self-administration will be important to establish causal roles for

SuM^VGluT2→MS neurons in nicotine reinforcement. While our data implicate β2-containing receptors, direct pharmacological or genetic manipulations will be required to test subunit-specific contributions.

The SuM has been extensively linked to arousal and stress (Pedersen et al., 2017; Escobedo et al., 2024; Zhang et al., 2026), and it projects to not only the medial septum but also the hippocampus, modulating oscillatory activity and functions such as spatial memory and anxiety (Luo et al., 2025). Notably, nicotine is recognized for its effects on memory and mood (Picciotto et al., 2000). While somewhat speculative, nicotine-induced activation of SuM neurons may impact not only nicotine’s reinforcing properties but also its effects on memory and mood. Thus, investigating whether nicotine’s established effects are mediated, at least in part, via the SuM represents a compelling avenue for future research.

The SuM may play a role in addiction beyond nicotine. Although this research found no significant changes in SuM neuron activity with cocaine or fentanyl, previous studies indicate that methamphetamine stimulates the SuM and increases local field potential (LFP). Suppressing these neurons reduces both methamphetamine-induced conditioned place preference (CPP) and locomotor sensitization (LS) (Ren et al., 2024). Melatonin, acting through MT2 receptors, can lessen methamphetamine-triggered SuM activation and related behaviors, suggesting that melatonin and similar receptor ligands could be valuable clinically for regulating SuM activity.

### Conclusion

Together, these findings identify the supramammillary region as a previously underappreciated hypothalamic substrate engaged by reinforcing doses of nicotine. By integrating molecular, behavioral, and in vivo physiological approaches, the present study links nicotinic receptor expression in SuM neurons with nicotine-regulated intake and rapid activation of SuM VGluT2 neurons during drug exposure. These results extend current models of nicotine reinforcement beyond canonical mesolimbic pathways and highlight hypothalamic–septal circuitry as a potential contributor to nicotine’s reinforcing effects.

### Ethical statement

The experimental protocol was approved by the Animal Care and Use Committee of the Intramural Research Program at NIDA and conducted in accordance with the Guide for the care and use of laboratory animals (National_Research_Council, 2011).

### Funding

This research was supported by the Intramural Research Program of the National Institute on Drug Abuse, National Institutes of Health (NIH). YA received a fellowship from Center on Compulsive Behaviors, Intramural Research Program, National Institutes of Health. The contributions of the NIH authors are considered Works of the United States Government. The findings and conclusions presented in this paper are those of the author(s) and do not necessarily reflect the views of the NIH or the U.S. Department of Health and Human Services. In addition, the research was supported by Smoking Research Foundation (Tokyo, Japan).

### Declaration of competing interest

The authors declare no conflict of interest.

## Acknowledgements

The authors thank Drs. C. D. Fowler, Z. X. Xi, and Y. Shaham and his group for advice on nicotine IVSA procedure. The authors also acknowledge the use of microscopic facility provided by the NIDA Microscope Core.

## Notes

### Competing Interest Statement

The authors have declared no competing interest.

## REFERENCES

1. Abizaid A, Liu ZW, Andrews ZB, Shanabrough M, Borok E, Elsworth JD, Roth RH, Sleeman MW, Picciotto MR, Tschop MH, Gao XB, Horvath TL (2006) Ghrelin modulates the activity and synaptic input organization of midbrain dopamine neurons while promoting appetite. J Clin Invest 116:3229–3239.

2. Cabeza de Vaca S, Carr KD (1998) Food restriction enhances the central rewarding effect of abused drugs. J Neurosci 18:7502–7510.

3. Caggiula AR, Donny EC, White AR, Chaudhri N, Booth S, Gharib MA, Hoffman A, Perkins KA, Sved AF (2001) Cue dependency of nicotine self-administration and smoking. Pharmacol Biochem Behav 70:515–530.

4. Carr KD (2007) Chronic food restriction: enhancing effects on drug reward and striatal cell signaling. Physiol Behav 91:459–472.

5. Carr KD, Tsimberg Y, Berman Y, Yamamoto N (2003) Evidence of increased dopamine receptor signaling in food-restricted rats. Neuroscience 119:1157–1167.

6. Carroll ME, France CP, Meisch RA (1979) Food deprivation increases oral and intravenous drug intake in rats. Science 205:319–321.

7. Contet C, Whisler KN, Jarrell H, Kenny PJ, Markou A (2010) Patterns of responding differentiate intravenous nicotine self-administration from responding for a visual stimulus in C57BL/6J mice. Psychopharmacology (Berl) 212:283–299.

8. Corrigall WA, Franklin KB, Coen KM, Clarke PB (1992) The mesolimbic dopaminergic system is implicated in the reinforcing effects of nicotine. Psychopharmacology (Berl) 107:285–289.

9. Dehkordi O, Rose JE, Asadi S, Manaye KF, Millis RM, Jayam-Trouth A (2015) Neuroanatomical circuitry mediating the sensory impact of nicotine in the central nervous system. J Neurosci Res 93:230–243.

10. Di Chiara G, Imperato A (1988) Drugs abused by humans preferentially increase synaptic dopamine concentrations in the mesolimbic system of freely moving rats. Proc Natl Acad Sci U S A 85:5274–5278.

11. Donny EC, Chaudhri N, Caggiula AR, Evans-Martin FF, Booth S, Gharib MA, Clements LA, Sved AF (2003) Operant responding for a visual reinforcer in rats is enhanced by noncontingent nicotine: implications for nicotine self-administration and reinforcement. Psychopharmacology (Berl) 169:68–76.

12. Donny EC, Caggiula AR, Mielke MM, Booth S, Gharib MA, Hoffman A, Maldovan V, Shupenko C, McCallum SE (1999) Nicotine self-administration in rats on a progressive ratio schedule of reinforcement. Psychopharmacology (Berl) 147:135–142.

13. Donny EC, Caggiula AR, Rowell PP, Gharib MA, Maldovan V, Booth S, Mielke MM, Hoffman A, McCallum S (2000) Nicotine self-administration in rats: estrous cycle effects, sex differences and nicotinic receptor binding. Psychopharmacology (Berl) 151:392–405.

14. Escobedo A, Holloway SA, Votoupal M, Cone AL, Skelton H, Legaria AA, Ndiokho I, Floyd T, Kravitz AV, Bruchas MR, Norris AJ (2024) Glutamatergic supramammillary nucleus neurons respond to threatening stressors and promote active coping. Elife 12.

15. Fowler CD, Kenny PJ (2011) Intravenous nicotine self-administration and cue-induced reinstatement in mice: effects of nicotine dose, rate of drug infusion and prior instrumental training. Neuropharmacology 61:687–698.

16. Ikemoto S, Qin M, Liu ZH (2006) Primary reinforcing effects of nicotine are triggered from multiple regions both inside and outside the ventral tegmental area. J Neurosci 26:723–730.

17. Imperato A, Mulas A, Di Chiara G (1986) Nicotine preferentially stimulates dopamine release in the limbic system of freely moving rats. Eur J Pharmacol 132:337–338.

18. Keller KL, Vollrath-Smith FR, Jafari M, Ikemoto S (2014) Synergistic interaction between caloric restriction and amphetamine in food-unrelated approach behavior of rats. Psychopharmacology (Berl) 231:825–840.

19. Kesner AJ, Shin R, Calva CB, Don RF, Junn S, Potter CT, Ramsey LA, Abou-Elnaga AF, Cover CG, Wang DV, Lu H, Yang Y, Ikemoto S (2021) Supramammillary neurons projecting to the septum regulate dopamine and motivation for environmental interaction in mice. Nat Commun 12:2811.

20. Kish GB (1955) Learning when the onset of illumination is used as the reinforcing stimulus. J Comp Physiol Psychol 48:261–264.

21. Le May MV, Hume C, Sabatier N, Schele E, Bake T, Bergstrom U, Menzies J, Dickson SL (2019) Activation of the rat hypothalamic supramammillary nucleus by food anticipation, food restriction or ghrelin administration. J Neuroendocrinol 31:e12676.

22. Lenoir M, Kiyatkin EA (2011) Critical role of peripheral actions of intravenous nicotine in mediating its central effects. Neuropsychopharmacology 36:2125–2138.

23. Leonardo M, Brunty S, Huffman J, Kastigar A, Dickson PE (2023) Intravenous fentanyl self-administration in male and female C57BL/6J and DBA/2J mice. Sci Rep 13:799.

24. Lindblom J, Johansson A, Holmgren A, Grandin E, Nedergard C, Fredriksson R, Schioth HB (2006) Increased mRNA levels of tyrosine hydroxylase and dopamine transporter in the VTA of male rats after chronic food restriction. The European journal of neuroscience 23:180–186.

25. Luo YJ et al. (2025) Segregated supramammillary-dentate gyrus circuits modulate cognitive and affective function in healthy and Alzheimer’s disease model mice. Neuron.

26. McGregor A, Roberts DCS (1993) Dopaminergic antagonism within the nucleus accumbens or the amygdala produces differential effects on intravenous cocaine self-administration under fixed and progressive ratio schedules of reinforcement. Brain Research 624:245–252.

27. National_Research_Council (2011) Guide for the care and use of laboratory animals, 8th Edition. Washington, D.C.: The National Research Academies Press.

28. Pedersen NP, Ferrari L, Venner A, Wang JL, Abbott SBG, Vujovic N, Arrigoni E, Saper CB, Fuller PM (2017) Supramammillary glutamate neurons are a key node of the arousal system. Nat Commun 8:1405.

29. Perkins KA, Donny E, Caggiula AR (1999) Sex differences in nicotine effects and self-administration: review of human and animal evidence. Nicotine Tob Res 1:301–315.

30. Picciotto MR, Kenny PJ (2021) Mechanisms of Nicotine Addiction. Cold Spring Harb Perspect Med 11:a039610.

31. Picciotto MR, Caldarone BJ, King SL, Zachariou V (2000) Nicotinic receptors in the brain. Links between molecular biology and behavior. Neuropsychopharmacology 22:451–465.

32. Picciotto MR, Zoli M, Rimondini R, Lena C, Marubio LM, Pich EM, Fuxe K, Changeux JP (1998) Acetylcholine receptors containing the beta2 subunit are involved in the reinforcing properties of nicotine. Nature 391:173–177.

33. Plaisier F, Hume C, Menzies J (2020) Neural connectivity between the hypothalamic supramammillary nucleus and appetite– and motivation-related regions of the rat brain. J Neuroendocrinol 32:e12829.

34. Pogun S, Yararbas G, Nesil T, Kanit L (2017) Sex differences in nicotine preference. J Neurosci Res 95:148–162.

35. Pontieri FE, Tanda G, Orzi F, Di Chiara G (1996) Effects of nicotine on the nucleus accumbens and similarity to those of addictive drugs. Nature 382:255–257.

36. Ren Q, Han W, Yue Y, Tang Y, Yue Q, Comai S, Sun J (2024) Melatonin Regulates Neuronal Synaptic Plasticity in the Supramammillary Nucleus and Attenuates Methamphetamine-Induced Conditioned Place Preference and Sensitization in Mice. J Pineal Res 76:e13006.

37. Rose JE, Corrigall WA (1997) Nicotine self-administration in animals and humans: similarities and differences. Psychopharmacology (Berl) 130:28–40.

38. Skibicka KP, Hansson C, Alvarez-Crespo M, Friberg PA, Dickson SL (2011) Ghrelin directly targets the ventral tegmental area to increase food motivation. Neuroscience 180:129–137.

39. Sorge RE, Clarke PBS (2011) Nicotine Self-Administration. In: Animal Models of Drug Addiction (Olmstead MC, ed), pp 101–132. Totowa, NJ: Humana Press.

40. Speakman JR, Mitchell SE (2011) Caloric restriction. Mol Aspects Med 32:159–221.

41. Stewart J, Hurwitz HMB (1958) Studies in light-reinforced behaviour III: The effects of continuous, zero and fixed-ratio reinforcement. Q J Exp Psychol 10:56–61.

42. Thomsen M, Caine SB (2006) Cocaine self-administration under fixed and progressive ratio schedules of reinforcement: comparison of C57BL/6J, 129X1/SvJ, and 129S6/SvEvTac inbred mice. Psychopharmacology (Berl) 184:145–154.

43. Thomsen M, Hall FS, Uhl GR, Caine SB (2009) Dramatically decreased cocaine self-administration in dopamine but not serotonin transporter knock-out mice. J Neurosci 29:1087–1092.

44. Walters CL, Brown S, Changeux JP, Martin B, Damaj MI (2006) The beta2 but not alpha7 subunit of the nicotinic acetylcholine receptor is required for nicotine-conditioned place preference in mice. Psychopharmacology (Berl) 184:339–344.

45. Wills L, Ables JL, Braunscheidel KM, Caligiuri SPB, Elayouby KS, Fillinger C, Ishikawa M, Moen JK, Kenny PJ (2022) Neurobiological Mechanisms of Nicotine Reward and Aversion. Pharmacol Rev 74:271–310.

46. Zhang J, Yu K, Zhang J, Chang Y, Sun X, Qian Z, Sun Z, Qiao Y, Liu Z, Ren W, Han J (2026) A stress-activated neuronal ensemble in the supramammillary nucleus produces anxiety-like behavior in male mice. Elife 14.

