## Supplemental Table for "Reinforcing Nicotine Doses Activate Supramammillary VGluT2 Neurons in Mice"

Supplementary Table 1. ANOVAs (SPSS v31.0.1.0)

| Figure  DV with test | Factor Name | Value | P Value | Partial Eta^2^ |
| --- | --- | --- | --- | --- |
| Fig. 2A  Nicotine Infusion with one-way repeated measures ANOVA | Dose (sphericity)  Dose (within) | 𝜒^2^ = 3.36; E_GG_ = 0.73  *F*_3, 15_ = 17.05 | = 0.656  < 0.001 | 0.77 |
| Fig. 2B  Leverpress with two-way repeated measures ANOVA | Dose (sphericity)  Lever X Dose (sphericity)  Lever (within)  Dose (within)  Lever X Dose | 𝜒^2^ = 8.32; E_GG_ = 0.49  𝜒^2^ = 21.90; E_GG_ = 0.35  *F_1_*_, 5_ = 12.71  *F*_3, 15_ = 12.77  *F*_1.01, 5.26_ = 1.94 | = 0.151  < 0.001  = 0.016  < 0.001  = 0.221 | 0.72  0.72  0.28 |
| Fig. 3A  Nicotine Infusion with two-way mixed measures ANOVA | Dose (sphericity)  Group (between)  Dose (within)  Group X Dose | 𝜒^2^ = 26.31; E_GG_ = 0.56  *F*_1, 22_ = 20.45  *F*_1.7, 36.7_ = 6.38  *F*_1.7, 36.7_ = 4.70 | < 0.001  < 0.001  = 0.006  = 0.020 | 0.48  0.23  0.18 |
| Fig. 3A  Nicotine Infusion with tests of within-subjects contrasts, two-way mixed measures ANOVA | Dose - Linear  Dose - Quadratic  Dose - Cubic  Group X Dose - Linear  Group X Dose - Quadratic  Group X Dose - Cubic | *F*_1, 22_ = 3.91  *F*_1, 22_ = 22.21  *F*_1, 22_ = 0.74  *F*_1, 22_ = 3.57  *F*_1, 22_ = 9.57  *F*_1, 22_ = 4.57 | = 0.061  < 0.001  = 0.399  = 0.072  = 0.005  = 0.044 | 0.15  0.50  0.03  0.14  0.30  0.17 |
| Fig. 3B  Nicotine group: Leverpress with two-way repeated measures ANOVA | Dose (sphericity)  Lever X Dose (sphericity)  Lever (within)  Dose (within)  Lever X Dose | 𝜒^2^ = 15.10; E_GG_ = 0.54  𝜒^2^ = 22.65; E_GG_ = 0.50  *F_1_*_, 13_ = 51.46  *F*_1.6, 21.1_ = 9.18  *F*_1.5, 19.6_ = 4.15 | = 0.01  < 0.001  < 0.001  = 0.002  = 0.041 | 0.80  0.41  0.24 |
| Fig. 3B  Saline control group: Leverpress with two-way repeated measures ANOVA | Dose (sphericity)  Lever X Dose (sphericity)  Lever (within)  Dose (within)  Lever X Dose | 𝜒^2^ = 133.71; E_GG_ = 0.38  𝜒^2^ = 51.08; E_GG_ = 0.34  *F*_1, 9_ = 0.73  *F*_1.1, 10.1_ = 0.24  *F*_1.0, 9.3_ = 0.06 | < 0.001  < 0.001  = 0.416  = 0.663  = 0.817 | 0.08  0.03  0.01 |
| Fig. 3C  Nicotine group: Infusion with two-way mixed measures ANOVA | Dose (sphericity)  Strain (between)  Dose (within)  Strain X Dose | 𝜒^2^ = 19.19; E_GG_ = 0.51  *F*_1, 12_ = 0.28  *F*_1.5, 18.2_ = 7.73  *F*_1.5, 18.2_ = 0.27 | = 0.002  = 0.61  = 0.006  = 0.708 | 0.02  0.39  0.02 |
| Fig. 3D  Nicotine group: Infusion with two-way mixed measures ANOVA | Dose (sphericity)  Sex (between)  Dose (within)  Sex X Dose | 𝜒^2^ = 20.89; E_GG_ = 0.56  *F*_1, 12_ = 0.11  *F*_1.4, 17.0_ = 8.21  *F*_1.4, 17.0_ = 0.74 | < 0.001  = 0.74  = 0.006  = 0.449 | 0.01  0.41  0.06 |
| Fig. 4C  Nicotine: AUC with one-way repeated measures ANOVA | Dose (sphericity)  Dose (within) | 𝜒^2^ = 12.48; E_GG_ = 0.47  *F*_1.4, 15_ = 9.34 | = 0.034  = 0.014 | 0.77 |
| Fig. 4C  Coc & fentanyl: AUC with one-way repeated measures ANOVA | Drug (sphericity)  Drug (within) | 𝜒^2^ = 1.10; E_GG_ = 0.77  *F*_2, 8_ = 2.78 | = 0.578  = 0.121 | 0.41 |

Abbreviations: DV, dependent variable; E_GG_, Greenhouse-Geisser epsilon value

Supplementary Table 2. Strain and sex of the nicotine and saline groups in the PR3 schedule experiment

| Strain | Sex | NIC | SAL |
| --- | --- | --- | --- |
| VGluT2 | Male | 5 | 4 |
|  | Female | 3 | 2 |
| C57BL/6 | Male | 3 | 3 |
|  | Female | 3 | 1 |
